# Reproductive resource allocation correlates with successful global invasion of a mosquito species

**DOI:** 10.1101/2024.07.18.604133

**Authors:** Ayda Khorramnejad, Claudia Alfaro, Stefano Quaranta, Alejandro Nabor Lozada-Chávez, Laila Gasmi, Hugo D. Perdomo, Laurent Roberto Chiarelli, Mariangela Bonizzoni

**Affiliations:** Department of Biology and Biotechnology, University of Pavia, Pavia, 27100, Italy

**Keywords:** Invasion, mosquitoes, reproduction, *Aedes albopictus*

## Abstract

The understanding of traits favoring biological invasions has been considered an essential step to predict which species would become successful invaders. Classical approaches test for differences between invasive vs. not invasive species and emphasize reproduction as a critical phenotype for successful establishment of an invasive species. However, cross-species comparisons underestimate intraspecies differences, which may be relevant in invasive species with highly genetically diverse populations. Here we capitalize on the well-characterized invasion history of the arboviral vector *Aedes albopictus*, which resulted in genetically-distinct native, old and invasive populations, and compared the reproductive capacity (fertility and fecundity), development (timing of egg development, oviposition patterns and egg hatching) and physiology (blood digestion and nutrient movement during oogenesis) across populations. We observed that invasive mosquitoes optimize their nutrient investment during development and oogenesis, which leads to increased egg production with respect to native and long adapted laboratory mosquitoes. This higher fecundity results from a delay in oogenesis and is accompanied by higher fertility. We further tested inheritance of reproductive traits via reciprocal crosses, which showed a higher fertility and fecundity in hybrids with respect to parental strains and a potential contribution of males to the reproductive success of invasive mosquitoes. Our results provide evidence that resource allocation during development and oogenesis influences the reproductive capacity of *Ae. albopictus* and manifests in population differences that correlate with their invasion success.

**Significance Statement:** In addition to being an essential process to ensure the survival of a species, reproduction is a key determinant for a species invasion success because it facilitates a species’ ability to establish in a new area. Reproduction is a complex phenotype that relies on intricate and timely interactions between genetic and physiological factors. Here we combined molecular, biochemical, and genetic approaches to show that efficient allocation of energetic resources during development and oogenesis fosters the reproductive output of *Ae. albopictus* mosquitoes and manifests as variation in the reproductive capacity of its geographic populations. These results are critical for predicting the invasion success of this species and tailoring effective control strategies.

## Introduction

Biological invasions are complex and multifaceted processes that have been recognized since the late 1950s as an increasing phenomenon with daunting ecological, sociological and economic consequences (1). Therefore, understanding of the key traits of invasive species has been considered an important step for the prediction of which species would become successful invaders, as well as for their management (2). A typical approach to identify such traits is the comparison of species with different invasion success (3–5). Experimental data obtained from many taxa using a cross-species comparison approach highlight the crucial role of the reproductive success of a species for its positive establishment in a new area, following introduction and preceding further spread (3). For example, invasive pine species tend to have a small seed mass, which is indicative, in these plants, of a larger seed number, high germinability and higher relative growth rate of seedlings, than non-invasive pine species (6). Data from invasive insects, birds, and vertebrate species adhere with life-history traits of invasive plants, with successful invasions having been associated with a species high reproductive capacity and short generation times (4, 7–9). However, the reproductive capacity of a species is a complex phenotype, which depends on its genetics, reproductive development and physiology. In many organisms, reproductive development and physiology are not static traits (10). Remarkably, there is limited research investigating whether populations of an invasive species differ in their reproductive development and physiology, resulting in population differences in their overall reproductive capacity that could influence their invasive success (9)

The Asian tiger mosquito *Aedes albopictus* (Skuse, 1894) is the archetype of a successful invasive species. Native of Asia, this mosquito colonized islands of the Indian Ocean and the Pacific, such as Hawaii and La Reunion Island, during the spice trade of the 17th-18th century (11). Through global increases in commerce and human travels during the past century, *Ae. albopictus* moved globally and has now established populations in every continent, except Antarctica (12). Based on this dual wave of colonization, current *Ae. albopictus* populations are designated native, old or invasive depending on their origin in Asia, the Indian Ocean and the Pacific, or elsewhere in the world (12,13). Genetic data from studies of the last decades consistently showed that classification of populations also reflects their genetic differentiation (13–15).

The reproductive capacity of *Aedes* spp. females is tightly coordinated with their nutritional status through hormones (16). Blood meal stimulates vitellogenesis by triggering the release of neuropeptides from neurosecretory cells in the mosquito brain (16–17). Additionally, acquisition of a blood meal and its digestion change the metabolic activity of the fat body resulting in the production of yolk precursor proteins, secretion of yolk proteins and lipids and their deposition in ovaries (18–20). Oviposition-related metabolism also promotes morphological and physiological changes in ovaries during oogenesis (20). *Aedes* spp. females emerge with two ovaries each divided into 50-60 ovarioles containing an oocyte, nurse cells and surrounding follicle cells forming the primary egg chamber (20). During vitellogenesis, developing oocytes grow synchronously through acquisition of proteins, lipids and carbohydrates to form mature eggs (20). The oocyte of the secondary egg chamber will develop into a mature egg after a secondary blood meal (20). Egg production is cyclical. One round of ovarian development from the time of blood intake to egg laying is called gonotrophic cycle (GC). In addition to influencing egg development, blood meal profound physiological effects are mediated by the induction or repression of specific genes (21) and are also the mechanism through which *Ae. albopictus* may acquire and later transmit pathogens, including the nematode *Dirofilaria immitis* and the arboviruses Dengue and Chikungunya, among the most public-health relevant ones (12). Because of this crucial role, behavioral, molecular and hormonal processes regulating, and being regulated by, a blood meal have been extensively studied in vector mosquitoes at the species level (22).

Here we shift our focus from species to populations and ask whether blood meal intake, its digestion and allocation of resulting resources differ among *Ae. albopictus* populations, impacting their overall reproductive capacity. We further studied ovarian physiology through micrographs, proteomic analyses and the potential inheritance of reproductive traits via reciprocal crosses. We observed that invasive mosquitoes optimize their nutrient investment during oogenesis, leading to increased egg production. This higher fecundity is associated with a delay in oogenesis and is followed by a higher fertility. Overall, our findings support the conclusion that resource allocation during development and oogenesis influences the reproductive capacity of *Ae. albopictus* populations, ultimately affecting their invasion success.

## Results

### Invasive mosquitoes have large bodies and high reproductive capacity

We established three laboratory populations from invasive *Ae. albopictus* mosquitoes sampled in Tapachula (Mexico), Crema and Pavia (Italy) (23,24). We also adapted to laboratory conditions eggs collected in the Indian Ocean Island of La Reunion. We measured the body size and the reproductive output (fecundity and fertility) of mosquitoes of these laboratory-adapted populations, within the first 10 generations of laboratory colonization, and compared them to mosquitoes of the Foshan (Fo) population (23).

We observed that mosquitoes derived from invasive populations, namely Tapachula (Tap), Crema (Cr) and Pavia (Pv), have wider wing length, in both sexes, and higher reproductive output compared to mosquitoes from the old population from La Reunion Island and Fo mosquitoes (Fig. 1, Panels A-C), but the reproductive output did not differ significantly among invasive mosquitoes (Table S1). Additionally, using whole genome sequencing data (WGS), we verified that our laboratory-adapted populations are genetically different, with levels of genetic diversity and pairwise Fst values comparable to those estimated across invasive, old and native *Ae. albopictus* populations (13,23,25) (Table S2 and S3).

**Figure 1.**
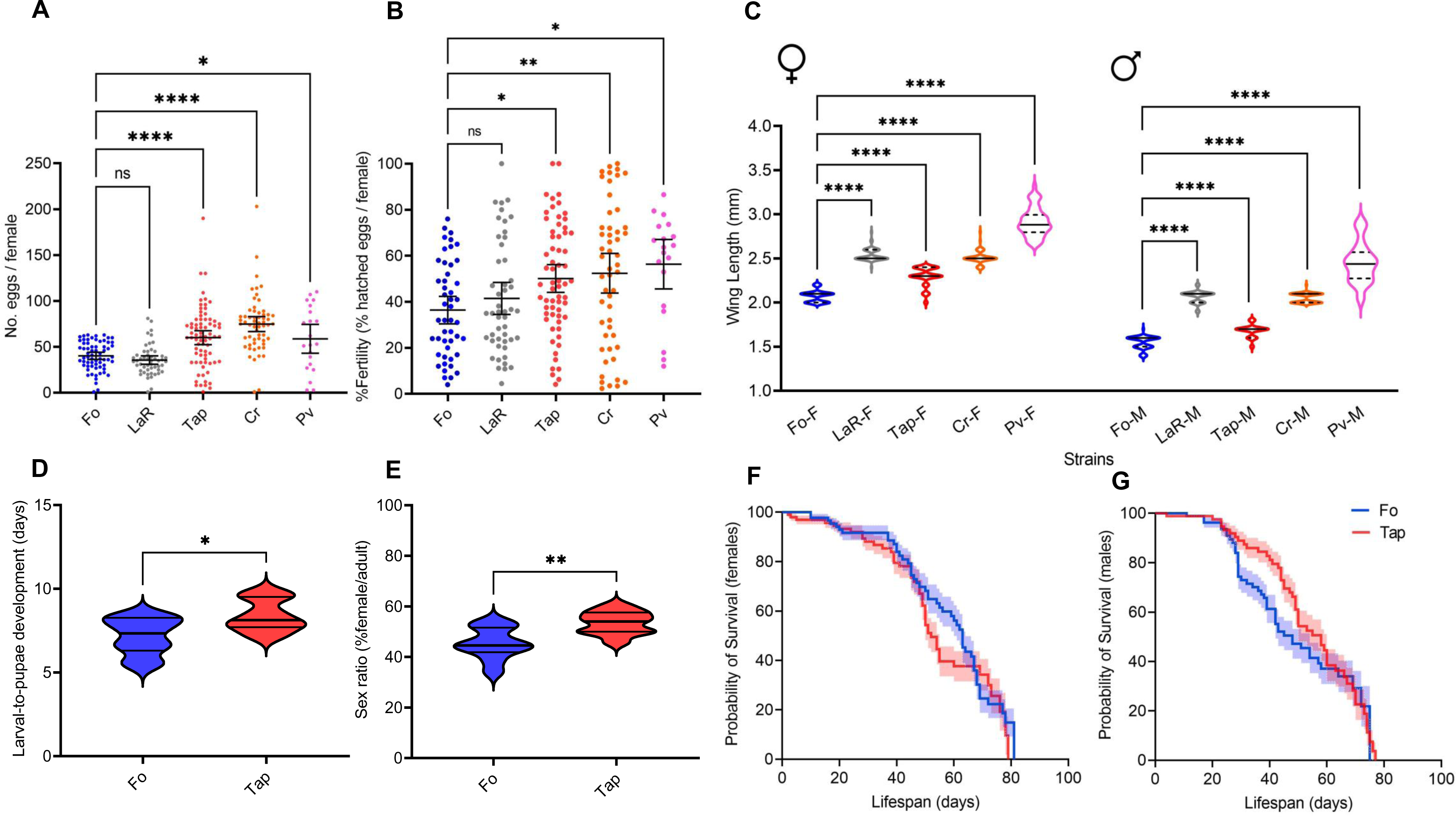
Phenotypic variations among *Aedes albopictus* populations. (**A**) Average number of eggs laid per female of the Foshan (Fo), La Reunion (LaR), Tapachula (Tap), Crema (Cr) and Pavia (Pv) laboratory-adapted populations (Fo, n=65; LaR, n=51; Tap, n=77; Cr, n=59; Pv, n=21). (**B**) Percentage of larvae developed from the eggs deposited by each female (Fo, n=48; LaR, n=49; Tap, n=62; Cr, n=52; Pv, n=20). (**C**) wing length (mm) of females and males; for all, at least 50 females and 50 males were analyzed. Differences were compared using a one-way ANOVA and error bars represent the 95% confidence intervals (**D**) Larval-to pupal developmental time (days). (**E**) Percentage of females in emerging adults (sex ratio). Differences were tested using an unpaired t-test. (**F**) Female and (**G**) male longevity. blue color represents Fo and red shows Tap mosquitoes. Differences were tested via Log-rank analyses. Ns represents not significant, * P-value<0.05, ** P-value<0.01, *** P-value<0.001, and **** P-value<0.0001. In all panels, data on Fo are in blue, LaR in gray, Tap in red, Cr in orange and Pv in pink.

### The higher reproductive capacity of Tap mosquitoes correlates with a longer developmental time

Among our invasive populations, Tap is more recent invasive population than Cr and Pv given that *Ae. albopictus* established in Italy in the 1990s but was first detected in tropical Chiapas in the early 2000 (12,26). On this basis, we selected Tap to further investigate the reproductive capacity of invasive mosquitoes. First, we extended the phenotypic comparison between Tap and Fo mosquitoes to their juvenile stages because developmental time can influence mosquito body size, which is directly correlated with a female reproductive capacity (Table S4) (27). We observed that larval-to-pupae developmental time was significantly faster in Fo compared to Tap mosquitoes (Fo=7.151±0.3422 days, Tap=8.395±0.3030 days, p value=0.0140) Fig. 1D) and that the larval viability was significantly higher in Tap compared to Fo (Tap=0.98±0.006, Fo=0.93±0.011, p value= 0.0020, Table S4). The sex ratio of emerging adults was different between populations, with significantly more females in Tap than Fo (Fig. 1E). We did not observe differences in adult longevity in either males or females between the two populations (Fig. 1 F,G).

We repeated assessment of Tap reproductive capacity after 31 generations in the laboratory and confirmed that in comparison to Fo, Tap mosquitoes have a higher fecundity (Tap=60.05 ± 3.79 eggs; Fo=38.70 ± 2.12 eggs, unpaired T-test, p value<0.0001) and fertility (Tap=50.10±3.01%, Fo=35.67±3%, p value=0.0018, Table S3) and that the higher reproductive output of Tap mosquitoes extends from the first (Fig. 2 A-B) to the second GC (Fig. 2 F-G; Table S4).

**Figure 2.**
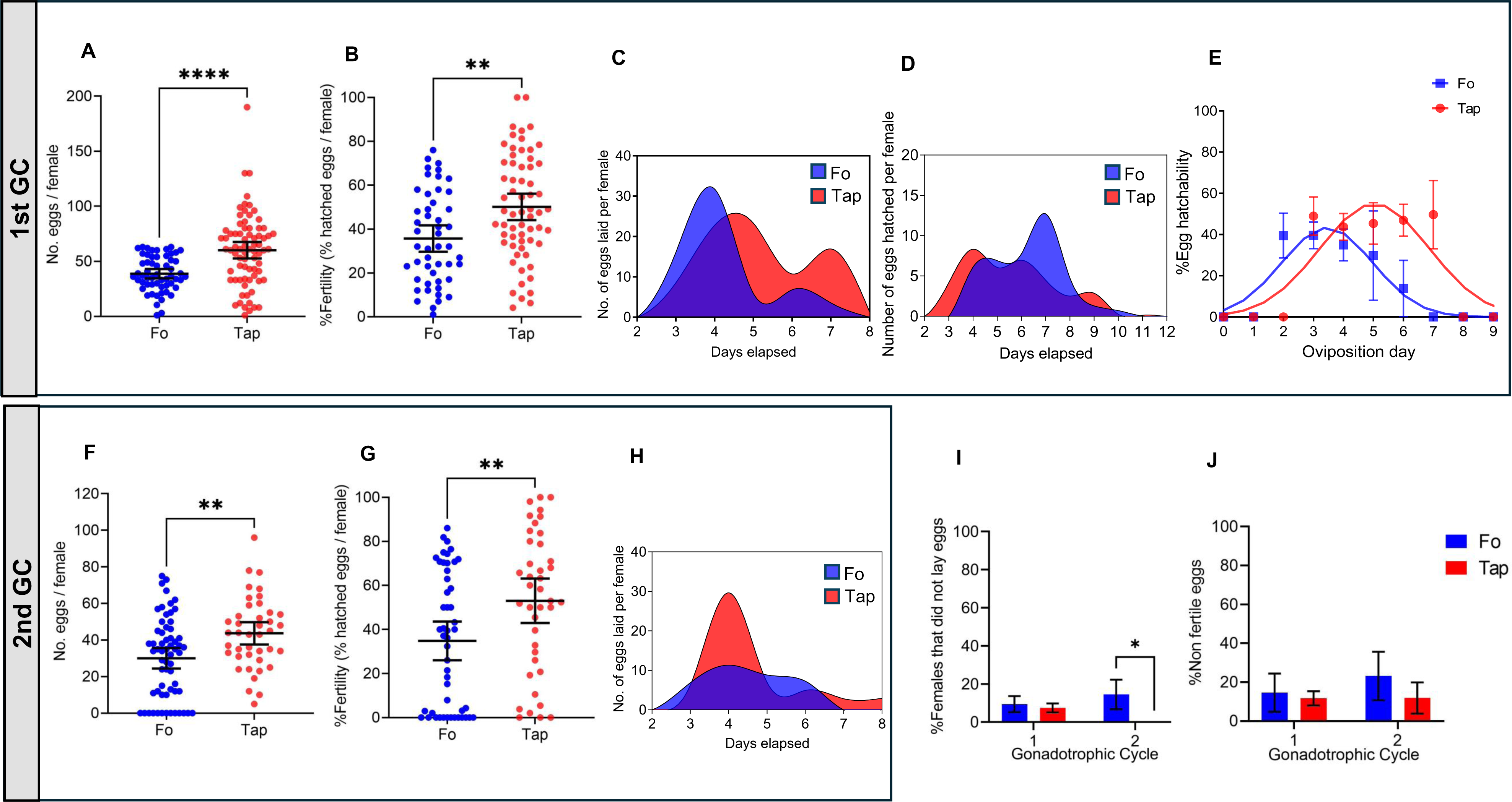
Reproductive output comparison between first and the second gonotrophic cycle (GC). **(A)** Fecundity of Tap (n=77) and Fo (n=57) females in the first GC. **(B)** Fertility of Tap (n=62) and Fo (n=49) females in the first GC. **(C)** Cubic spline curves showing the oviposition patterns of Fo and Tap mosquitoes during the first GC. Cubic spline curves were generated by interpolating the number of eggs laid by Fo and Tap females from 2 to 8 days post blood meal (pBM). (**D**) Cumulative pattern of egg hatching during the first GC (n=44 and n=49, for Fo and Tap, respectively). (**E**) Gaussian bell-shaped distribution shows viability of each batch of eggs deposited by Fo and Tap females in relation to their deposition day during the first GC. **(F)** Fecundity of Tap (n=41) and Fo (n=62) females in the second GC. **(G)** Fertility of Tap (n=40) and Fo (n=49) females in the second GC. (**H**) Cubic spline curves showing the oviposition patterns of Fo (n=28) and Tap (n=11) mosquitoes during the second GC. Cubic spline curves were generated to interpolate eggs laid by Fo and Tap females from 2 to 8 days PBM. (**I**) Percentage of females that did not lay eggs during the first (1) and second (2) GC. (**J**) Percentage of laid eggs, which did not hatch, during the first (1) and second (2) GC. For (A), (B), (F), (G), (I), (J), differences were tested using an unpaired t-test. Ns represents not significant, ** P-value<0.01 and **** P-value<0.0001. In all panels, data of Fo and Tap are in blue and red, respectively.

### Tap mosquitoes show a delay in oviposition in the first gonotrophic cycle

We tested whether the higher reproductive capacity of Tap with respect to Fo mosquitoes is linked to their pattern of oviposition by counting the number of eggs deposited daily from the second to the eighth day post blood meal (PBM), both during the first (Fig. 2C) and the second GCs (Fig. 2H; Table S4). The oviposition curves of Fo and Tap mosquitoes were significantly different in both GCs (first GC: F value=13,61; second GC: F-value=6,167). During the first GC, the number of eggs laid daily by Fo females peaked between three to four days PBM and then gradually declined. More than 95% of Fo females deposited their eggs in one or two batches, each with an average clutch size of 20.38 (±2.99) eggs (Fig. 2C, Table S4); less than 2% of tested females laid more than two clutches of eggs throughout the six days of our observation. In contrast, there were two peaks of oviposition for Tap females, one between four and five days pBM and a second one seven days pBM; 21.12% (±9.47) of Tap mosquitoes laid more than two clutches of the eggs, with an average clutch size of 22.80 (±0.54) eggs (Fig. 2C).

We further tested whether the observed differences in the oviposition pattern between Fo and Tap females extend to egg hatchability by individually assessing the fertility of females from the second to the eighth day pBM. We observed that the first batch of eggs deposited by Tap females hatched faster than that produced by Fo females (Fig. 2D). We further saw that the hatchability of Fo eggs peaked two days post oviposition (PO) (38.63%±6.43), then gradually declined, reaching a minimum, of 13.85±7.88% hatchability six days PO, which was significantly lower than the maximum hatchability value (p value<0.05). Viability of Tap eggs remained constant independently of their oviposition day (Fig. 2E, Table S4). During the second GC, oviposition patter of Fo mosquitoes showed no peak (Fig. 2H). Tap mosquitoes started laying eggs earlier pBM in the second GC, with an oviposition peak two days pBM (Fig. 2H). We further observed that while no significant differences in the number of sterile females was detected in the first GC (p value>0.05) (9.45%±2.42 in Fo *vs.* 7.41%±1.33 in Tap), the percentage of females laying no eggs increased to 14.51% (±4.48) in Fo, but decreased to zero in Tap mosquitoes during the second GC (Fig. 2 I-J; Table S3). Overall, these results show that Tap females have a delay in oviposition with respect to Fo during the first GC; this delay is accompanied by an overall higher fecundity and fertility and the reproductive advantage of Tap *vs.* Fo mosquitoes continues in the second GC.

### Oviposition delay in Tap females has physiological bases

The observed delay in the oviposition pattern of Tap females after the first GC could be a behavioral trait whereby mosquitoes retain mature eggs in their ovaries, or it could result from the physiology of egg production. We combined two approaches to discriminate between these alternatives. First, we studied egg development in maturing ovaries by acquiring micrographs at intervals between 6 to 72 hours (h) pBM starting at 24 hpBM (Fig. 3A). Second, we compared temporal changes in ovarian proteins between Fo and Tap females (Fig. 3 B-D).

**Figure 3.**
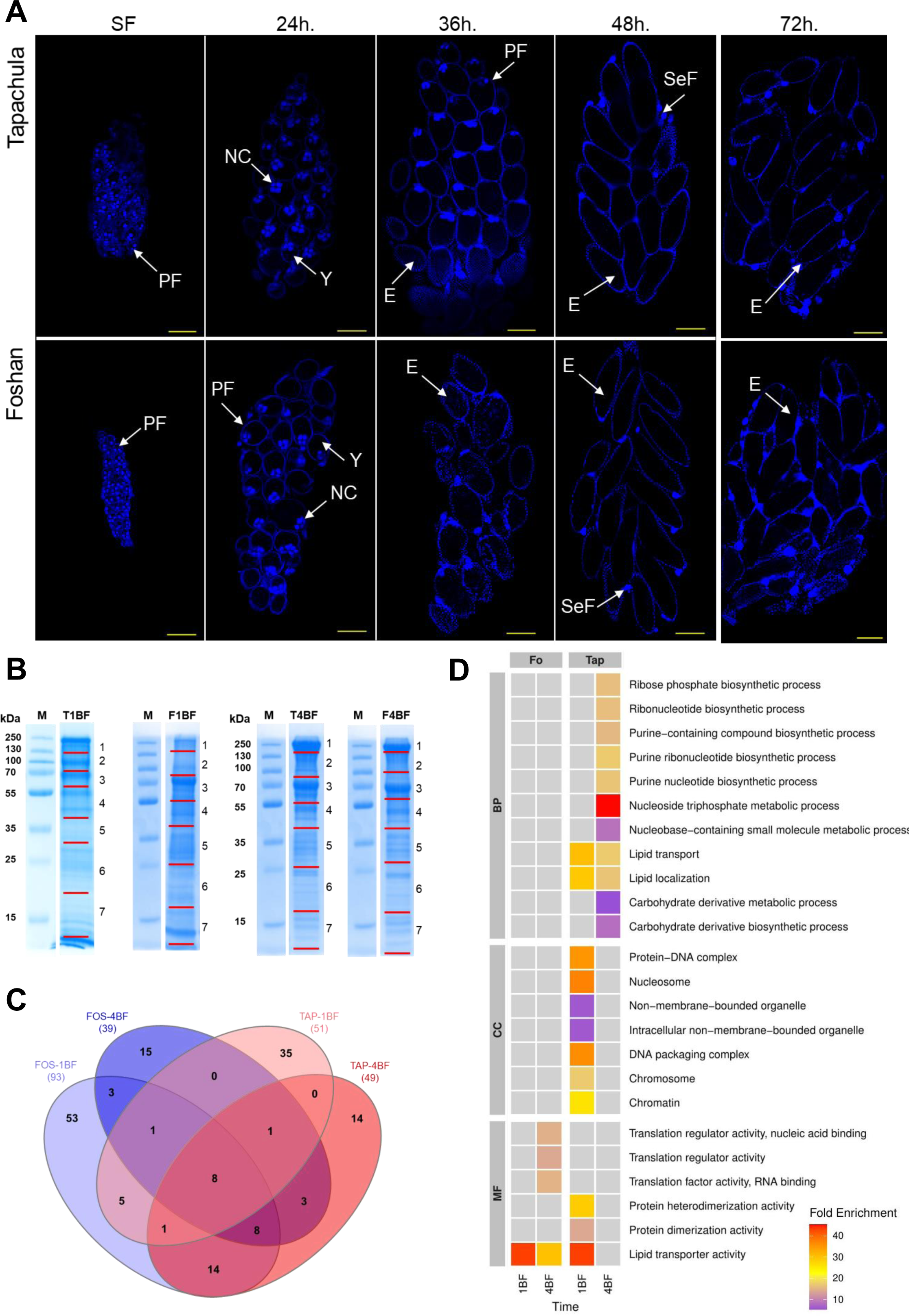
Morphology and proteomics of ovaries. (**A**) Micrographs of ovaries of females of the Fo and Tap populations sampled sugar-fed (SF), 24, 36, 48 hours pBM. The nuclei were visualized using 4’, 6-diamidino-2-phenylindole (DAPI) staining. Arrows point to primary follicles (PF), nurse cells (NC), yolk (Y), eggs (E) and secondary follicles (SeF). The yellow solid line represents a scale of 50 µm. (**B**) The SDS-PAGE analysis of ovarian proteins extracted from of ovaries of Tap (T) and Fo (F) females dissected one (1) and four (4) days pBM. Lanes M correspond to the PAGERuler Plus Prestained Protein Ladder (ThermoFisher Scientific). Red lines mark the gel slices that were excised. (**C**) Venn diagram showing the number of proteins identified by LC/MS in each condition/population. (**D**) Functional enrichment of proteins from ovaries of Fo and Tap mosquitoes sampled one and four days PBM. MF stands for molecular functions, CC for cellular constituents and BP for biological processes.

We captured images of ovaries of Fo and Tap females 24, 30, 36, 48 and 72 hpBM and analyzed them in comparison to images of ovaries of sugar-fed (SF) females (Fig. 3A). We saw primary follicles with little or no yolk in all SF females (Fig. 3A). We observed primary follicles with nurse cells and an increasing amount of yolk in ovaries of both Fo and Tap females sampled 24 and 30 hpBM. At 36 hpBM, we saw a clear difference in the images of Fo and Tap mosquitoes (Fig. 3A). We observed mature eggs and few primary follicles in ovaries of Fo females; whereas ovarioles containing mostly primary follicles with high amounts of yolk were seen in Tap females. Mature eggs in the ovaries of both populations were seen after 48h pBM. These results support the conclusion that the observed delay in oviposition of Tap with respect to Fo females is caused by different dynamics of egg formation and not by egg retention in Tap mosquitoes.

We further compared the protein profile of ovaries of fully-engorged blood-fed Tap and Fo mosquitoes, one- and four-days pBM, which represent the trophic and post-trophic stages of vitellogenesis in mosquitoes, respectively (16,20) The quantity of total proteins extracted from ovaries one day pBM did not differ between Fo and Tap mosquitoes (47.96±4.99 µg/mg in Fo and 42.08±6.93 µg/mg in Tap), but the complexity and abundance of proteins were different (Fig. 3B). A different trend was seen four days pBM when overall protein quantity was higher in Tap vs Fo ovaries (total protein content was 107.72±17.65 µg/mg in Fo and 163.92±18.88 µg/mg in Tap, p value=0.0110, Fig. S1), but the profile of proteins was similar. Tap ovaries sampled one day pBM were enriched in proteins associated with carbohydrate metabolic process and proteins linked to non-membrane bounded organelle, which are associated with cell structure and motility (28) (Fig. 3C-D, Table 1, Table S5). At the same time point pBM, proteins found uniquely in Fo ovaries were enriched in lipid transporter activity and proteins associated with oocyte and egg development. A total of 15 proteins (Fig. 3C, Table 1), among which is notable the presence of vitellogenin precursor (AALF008766), (Table S5), were found in ovaries of both Fo and Tap females one day PBM (Fig. 3 C-D). Four days pBM, we detected in Tap ovaries proteins enriched in functions such as carbohydrate metabolic processes, proteolysis, and lipid localization related to oocyte development (29). At the same time point, proteins exclusive of Fo ovaries were enriched in proteins linked to translation regulatory activity, lipid transporter and proteins that mediate the late stages of oogenesis and egg activation (30). A total of 20 proteins, some of which are involved in oogenesis (e.g. vitellogenin precursor [AALF008766]) were found in ovaries of both Fo and Tap mosquitoes four days pBM (Table 1). Based on the observed delay in oviposition in Tap mosquitoes with respect to Fo ones, we compared the profiles of proteins detected in ovaries sampled four days pBM in Tap and one day pBM in Fo females (Fig. 3C, Table 1). We found 31 proteins associated with oogenesis, carbohydrate metabolic process, protein biosynthesis and glycolytic process). Importantly, none of these 31 proteins was found in ovaries of Tap mosquitoes sampled one day pBM, except for vitellogenin-A1 precursors (AALF008766 and AAEL010434) (Table S5). Altogether, proteomics data support a delay in the biochemical activity of Tap mosquitoes for oogenesis aligning with fitness results and data from ovarian micrographs.

**Table 1.**
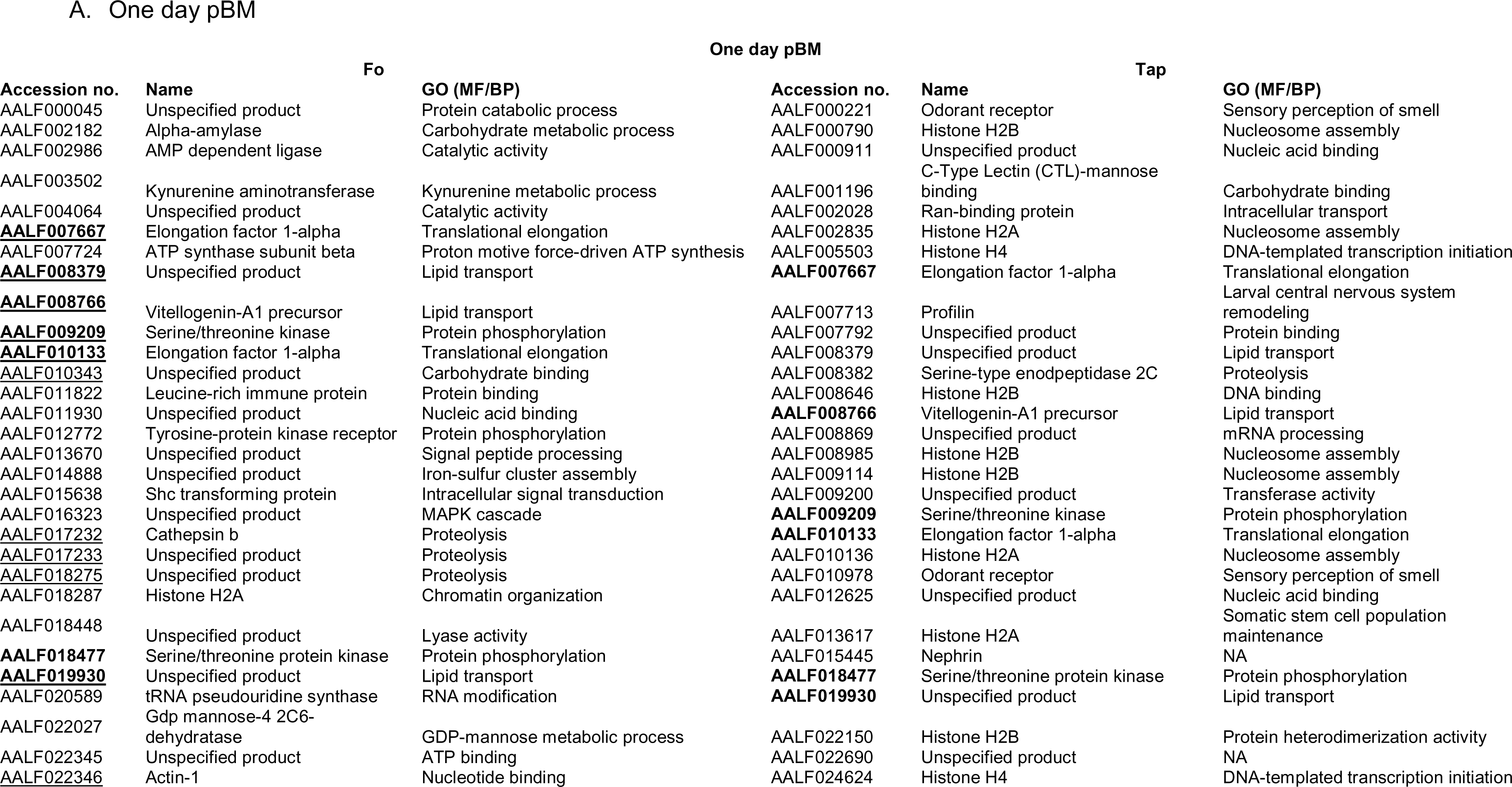
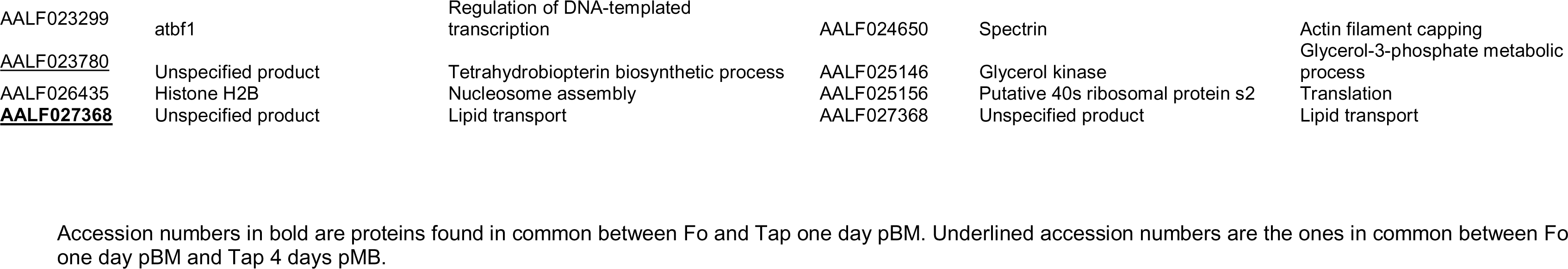
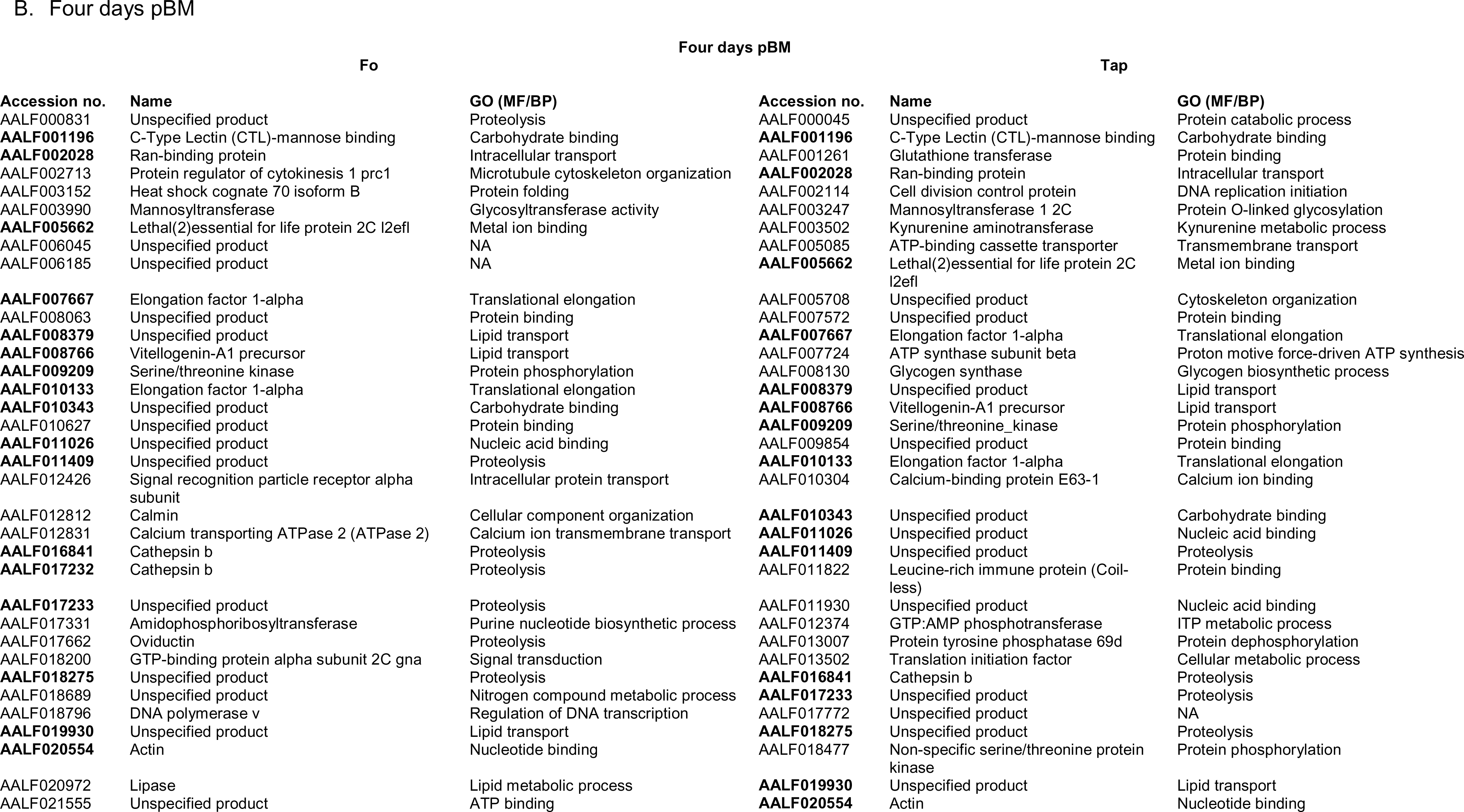
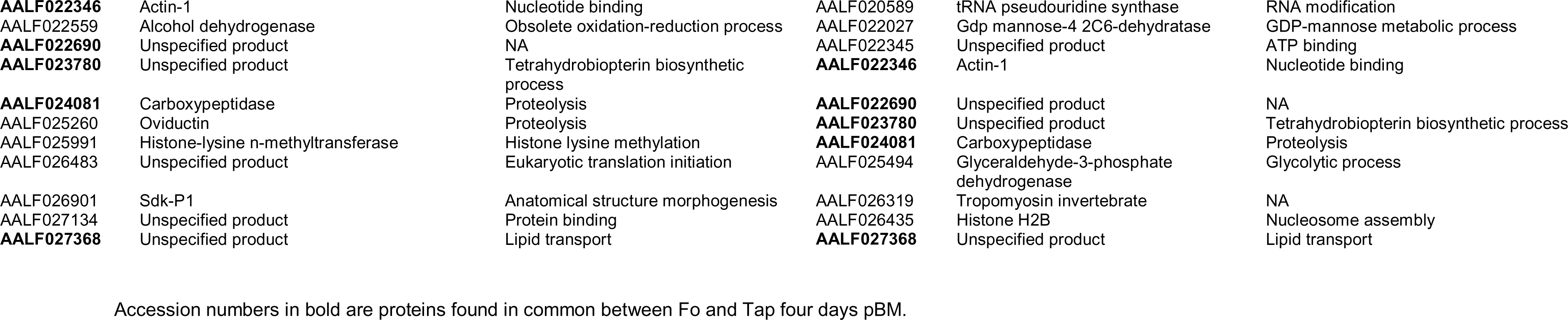
List of the identified proteins by LC-MS/MS analysis in ovaries of Fo and Tap one (A) and four (B) days pBM with their gene ontology based on either molecular function (MF) or biological process (BP).

### Tap mosquitoes efficiently allocate resources during oogenesis

Following acquisition and digestion of a blood meal, yolk precursor proteins are produced in the fat body, secreted into the hemolymph and transported to oocytes for egg production (20). To test if the delay in oogenesis we observed in Tap *vs.* Fo females depends on oviposition-related metabolism, we studied blood meal intake and we quantified trypsin-like enzyme activity, as a proxy for blood protein digestion (16); we further measured comparatively the accumulation and depletion of energy storage in the fat body and the absorption of lipids and proteins in the ovaries (Fig. 4).

**Figure 4.**
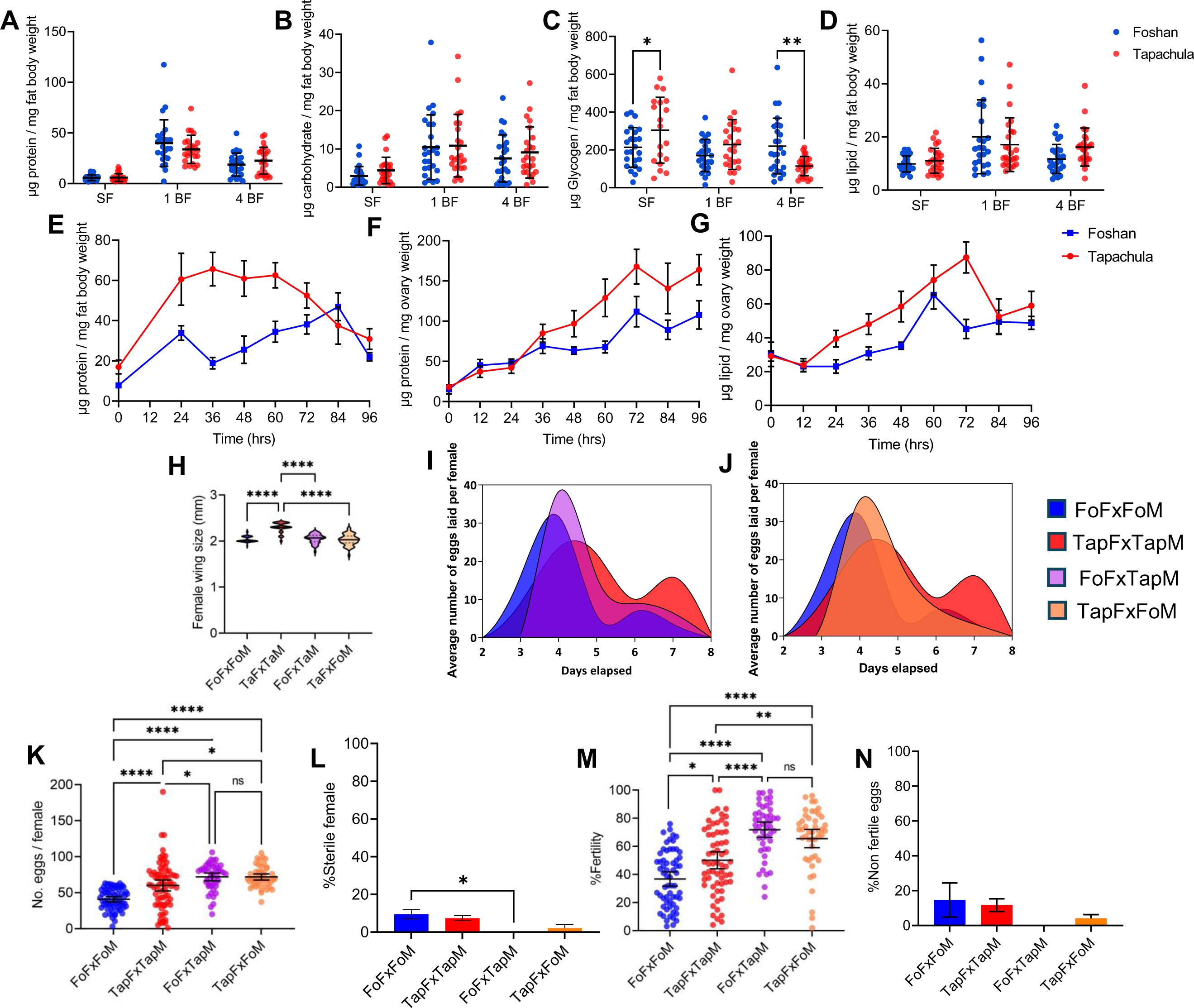
Oviposition-related metabolism and inheritance of reproductive traits. **(A)** Protein content in the fat body of Tap (n=24, for each timepoint) and Fo (n=25, for each timepoint) females before a blood meal (SF), 1 and 4 days pBM. (**B**) Carbohydrate content in the fat body of Tap (n=24, for each timepoint) and Fo (n=25, for each timepoint) females before a blood meal (SF), 1 and 4 days pBM. (**C**) Glycogen content in the fat body of Tap (n=24, for each timepoint) and Fo (n=25, for each timepoint) females before a blood meal (SF), 1 and 4 days pBM. (**D**) Lipid content in the fat body of Tap (n=24, for each timepoint) and Fo (n=25, for each timepoint) females before a blood meal (SF), 1 and 4 days pBM. (**E**) Protein content in the fat body of females before blood feeding (time 0) and 24, 36, 48, 60, 72, 84, and 96 h pBM. (**F**) Protein content in ovaries of Fo and Tap females before blood feeding (time 0) and 24, 36, 48, 60, 72, 84, and 96 h pBM. (**G**) Protein content in ovaries of Fo and Tap females before blood feeding (time 0) and 24, 36, 48, 60, 72, 84, and 96 h pBM. (**H**) Wing size of F1 progeny of parental Fo (n=50) and Tap (n=50) crosses and reciprocal crosses of Fo females and Tap males (n=38) or Fo males and Tap females (n=47), respectively. (**I**) Oviposition patterns of F1 progenies of parental Fo (n=49) and Tap (n=49) crosses and the reciprocal cross of Fo females and Tap males (n=48). (**J**) Oviposition patterns of F1 progenies of parental Fo (n=49) and Tap (n=49) crosses and the reciprocal cross of Fo males and Tap females (n=48). (**K**) Fecundity of F1 progeny of parental Fo (n=64) and Tap (n=75) crosses and the reciprocal crosses of Fo females and Tap males (n=83) or Fo males and Tap females (n=47). (**L**) Percentage of sterile females among the F1 progeny of parental Fo and Tap crosses and the reciprocal crosses of Fo females and Tap males (purple) or Fo males and Tap females (orange) (**M**) %Fertility of F1 progeny of parental Fo (n=63) and Tap (n=62) crosses and the reciprocal crosses of Fo females and Tap males (n=47) or Fo males and Tap females (n=46) (**N**) Percentage of laid eggs that did not hatch among the F1 progeny of parental Fo and Tap crosses and the reciprocal crosses of Fo females and Tap males or Fo males and Tap females. In panels, (**A**)-(**D**), (**K**) and (**M**), error bars represent the 95% confidence intervals. In panels, (**E**)-(**G**), (**L**) and (**N**), the error bars represent the SEM. A two-way ANOVA was used to analyze panels (**A**)-(**H**) and (**K**)-(**N**). In all plots, * stands for p value<0.05, ** p value<0.01, *** p value<0.001, and **** p value<0.0001. ns stands for no statistical significance.

We weighed 3-5-day old mosquitoes and then offered them a blood meal; after mosquitoes had flown to a resting spot, we weighed them again to avoid having quantification of blood intake biased by diuresis (Fig. S2). Blood intake was statistically higher in Fo than Tap females (p value=0.0028), with Fo females gaining 1.517±0.026 mg and Tap 1.052±0.066 mg after a blood meal (Fig. S2). Protease activity of trypsin-like enzymes showed similar trends in Fo and Tap mosquitoes (Fig. S2). We detected protease activity from 6 to 60 hpBM, with a peak at 24 hpBM in both Fo and Tap females. We registered a significantly higher activity of trypsin-like enzymes in Tap *vs.* Fo mosquitoes 24 hpBM (p value=0.0497). These results suggest that Tap mosquitoes process blood meal more efficiently than Fo. To confirm this result, we compared the nutritional status of SF mosquitoes with that of mosquitoes one- and four-days pBM by measuring the content in protein, carbohydrate, glycogen and lipid of their fat bodies. We observed that the trends in accumulation and depletion of all biochemicals was similar in both populations (Fig. 4A-D) with higher levels of tested nutrients in the fat body 24h after blood meal than in the pre-vitellogenic stage. Four days pBM we observed a reduction in the content of proteins, carbohydrates, and lipids in the fat body of both Tap and Fo mosquitoes (Fig. 4 A-B,D). We also observed that Tap mosquitoes deplete glycogen reserves quicker than Fo following the blood meal, although Tap females were more enriched in glycogen content before blood meal (Fig. 4C).

We further tracked the mobilization of proteins from the fat body and the accumulation of proteins and lipids in the ovaries at 12h intervals pBM (Fig. 4 E-G). We saw that the accumulation of protein in the fat body of Tap females gradually increased up to 60 hpBM, then gradually declined and reached its minimum level 96 hpBM. In contrast protein content in the fat body of Fo females dropped 36 hpBM and then gradually increased reaching its maximum level 84 hpBM, to decline again 96 hpBM (Fig. 4E). Protein mobilization from fat body to ovary showed a similar pattern in both Tap and Fo mosquitoes, with a gradual increase and a peak at 72 hpBM (Fig. 4F). Content of lipids in ovaries reached a peak 60 or 72 h pBM in Fo or Tap females, respectively (Fig. 4G). Together, these results demonstrate that Tap females better synchronize protein shifts from the fat body to ovaries pBM than Fo. Moreover, the trend in lipid content in ovaries aligns with the observed delay in egg production in Tap females.

### Cross-mating alters reproductive traits

We tested the genetic inheritance of fitness and reproductive traits by performing reciprocal crosses between Fo and Tap mosquitoes and studied wing length, fecundity, fertility and oviposition patterns of the F1 progeny (Fig. 4H-N). Wing length and oviposition patterns of the F1 progeny of both reciprocal crosses showed intermediate phenotypes with respect to those observed in parental populations suggesting Mendelian inheritance of quantitative traits (Fig. 4H-J). In contrast, fecundity and fertility values were higher in the F1 progeny of both reciprocal crosses than in either Tap and Fo parental mosquitoes (Fig. 4, K and M; Table S6). We further observed that the percentage of sterile females was significantly lower in the F1 progeny of crosses between Fo females and Tap males with respect to crosses of parental Fo, but not parental Tap mosquitoes (Fig. 4); the same trend was observed for the viability of eggs (Fig. 4N). These results suggest a contribution of Tap males in modulating the reproductive output of Fo females.

## Discussion

The arboviral vector *Ae. albopictus* is an aggressive invasive species that moved globally in less than 60 years (12). The success of this rapid invasion has been attributed mostly to its ability to produce desiccation-resistant eggs and undergo photoperiodic diapause (12,31). However, the native range of *Ae. albopictus* includes both temperate and tropical regions and native populations are under continuous gene-flow (32). Additionally, while diapause could have favored overwinter survival, establishment of *Ae. albopictus* populations in a new environment requires population growth over several generations. By combining fitness, physiological, proteomics and genetic data, we show here that invasive *Ae. albopictus* mosquitoes optimize their nutrient investment during development and following a blood meal, leading to a delay in oogenesis, but resulting in an increased egg production and higher fertility with respect to old and long-laboratory adapted mosquitoes.

Population differences in the reproductive capacity of *Ae. albopictus* have both physiological and genetic bases. Maintained under the same conditions and offered nutrition *ad libitum*, invasive Tap mosquitoes showed a longer developmental time, which correlated with both a larger body size and a higher content of glycogen 3-5 days post eclosion than Fo mosquitoes. Prolonged larval developmental time in Tap mosquitoes was also accompanied by sex ratio distortion with Fo having a higher percentage of males in emerging adults. Seasonal changes, day length, larval density in rearing containers, sex-specific responses to hatching stimuli and sex-selective predation can result in sex-ratio imbalances (33,34). Here, the observed sex ratio distortion could be due to lower mortality of males than females in Fo either as embryos and/or as larvae, as both fertility and larval viability were significantly lower in Fo than Tap mosquitoes.

While the positive correlation between female size and fecundity has been demonstrated previously in insects (35), we show that higher fecundity also depends on efficient resource allocation after a blood meal, which entails a delay in oogenesis. Despite blood meal intake being lower in Tap than Fo, fat body and ovaries of Tap were more enriched in protein, indicating that Tap females optimized blood meal digestion to allocate resources for egg production. In ovaries of Tap females, augmentation of lipids coordinated with protein accumulation, while lipid flux and protein deposition into developing ovaries did not follow a similar trend in Fo females. Tap females had lipids in ovaries later than Fo resulting in a delay in oogenesis and egg deposition. However, more efficient acquisition of nutrients allowed Tap females to sustain egg production longer than Fo, resulting in higher fecundity and fertility. The observed delayed in oogenesis of Tap mosquitoes indicates that maximizing allotment in the first clutch of eggs is associated with substantial benefits. Indeed, life-history theory predicts that early reproduction is more likely to be successful, and therefore more valuable, than later reproduction because the chance of mortality increases with age (36). We cannot exclude that shift of nutrients from the blood meal to reproduction may result in a fitness cost in terms of longevity. However, when we tested longevity and protein content in non blood-fed females we did not observe significant differences between Fo and Tap. This result is probably related to the fact that mosquitoes were kept at optimal rearing condition with unlimited access to nutrients. In Tap mosquitoes, optimal resource allocation translated into a delay in oogenesis and in the timing of oviposition but resulted in an overall higher fertility. This result is highly significant considering that Tap is a recent invasive population (12,26) and supports the conclusion that resource allocation during development and oogenesis influences the reproductive capacity of *Ae. albopictus,* favoring its invasion success.

We further show that Tap and Fo mosquitoes are genetically different and in reciprocal crosses female size was intermediate between that of parental strains, suggesting Mendelian inheritance of quantitative traits, hybrids displayed superior fecundity and fertility over their parents. Hybrid vigor, or heterosis, is a well-known phenomenon in agricultural settings (37) and is also beginning to be recognized among insects (38). The detection of heterosis is important for *Ae. albopictus* because its invasive populations have been shown to be genetically mixed due to human-mediated transport of eggs and overlaying introductions from both tropical and temperate areas (12–15). This invasion dynamic and our results on the vigor of the hybrids support the hypothesis that observed population differences in the reproductive capacity are adaptive. In addition to being facilitated by intraspecific hybridization in the introduced range, which may have created novel genotypes, rapid adaptation in *Ae. albopictus* reproductive capacity could be induced by severe changes in the selection regime imposed by new environments (38). The hypothesis that *Ae. albopictus* can undergo rapid adaptation is supported by results on the distribution of mutations predictive of pyrethroid-resistance, which appeared rapidly and independently across different populations (39). Moreover, there are frequent examples of invasive species adapting quickly to new environments. A well-studied example is the rapid evolution of *Drosophila suboscura* in the New World, which was demonstrated by the detection of rapidly evolving concordant patters of chromosomal inversions across latitudinal clines in three different continents (40). As an alternative explanation to the hypothesis of rapid adaptation, *Ae. albopictus* invasive populations may have a high reproductive capacity because they derive from regions prone to ecological disturbance. It has been hypothesized that fluctuating environments might favor organismal flexibility or the selection for evolvability by accumulation and maintenance of genetic variation (41). Indeed, recent population genomic studies showed that *Ae. albopictus* invasive and native populations retain similar levels of genomic diversity and that invasive populations can become sources of highly-adaptable founder strains promoting further invasions (13,25).

Analysis of more invasive populations sampled from different latitudes and environments is necessary to understand the origin and causes of the observed differences in the reproductive capacity of *Ae. albopictus* populations. However, our results clearly show that this difference has both genetic and physiological basis and is an important determinant of the species invasion success in addition to photoperiodic diapause.

## Materials and Methods

### Mosquito

We used mosquitoes of the long-established laboratory Fo population (23), along with mosquitoes adapted to the laboratory from eggs collected between 2017 and 2022 from La Reunion Island, Tapachula, Crema and Pavia (23–24). Briefly, eggs were collected from multiple ovitraps across each site or from adults (23–24). For the Pv laboratory-adapted population, adult mosquitoes were sampled in the summer of 2022 in numbers to generate at least 100 adults for the G1 laboratory generation. We reared all these five populations in parallel in Binder KBWF climatic chambers under constant conditions (28°C ± 1°C; 70-80% relative humidity; photoperiod of 12h:12h light:dark). Larvae were reared in plastic containers (BugDorm 19×19×6 cm) at a controlled density (around 200 larvae in 1 L of water) to prevent competition. We provided food daily in the form of fish food (Tetra Goldfish Gold Colour). Adults were kept in 30 cm^3^ cages and fed using cotton soaked with 20% sugar. Adult females were given commercial defibrinated mutton blood (Biolife Italiana) using a Hemotek blood membrane feeding apparatus.

### Fitness assessment

Fitness assessment was conducted within the first 10 generations after colony establishment for Tap, Cr, Pv and LaR mosquitoes. Fitness assessment was repeated at generation 31 for the Tap laboratory-adapted population. For each population, three replicates of 100 eggs each were hatched in plastic containers (17×6.5×12 cm) with 200 mL of water and reared at the standard conditions described above. Larval to pupal developmental time was recorded until pupation. Emerged adults were sexed. A sample of 50 females and 50 males was collected for wing length measurement as a proxy for body size, as previously described (24). For each population, a total of 200 females and 200 males were separated at emergence and the survival of every individual was monitored until death. We also monitored oviposition pattern, fecundity and fertility. Briefly, at least 50, 3-5 days old, females were placed in cages with 50 males to mate. After 7 days, females were offered a blood meal and we collected into individual circular cups only fully engorged females. We provided a damp filter paper for egg deposition two days after BM. We monitored the number of eggs laid by each female for the following 6 days to determine their oviposition pattern. We also measured the total number of eggs laid per female (fecundity). Next, we dried each filter paper for two days, before placing them in water for egg hatching and recording the number of larvae per female (fertility). In parallel, we took a minimum of 30, 3-5 days old, mated females per population and weighed them on a microbalance (Mettler AC100) before and after blood meal. The difference in weight corresponded to the amount of blood ingested (blood intake).

We used Prism 9.1 (GraphPad) for statistical analyses of life-history traits. We verified the normality of each dataset with the Shapiro-Wilk test (42) and proceeded accordingly. We tested differences in wing length, fecundity and fertility among the different laboratory-adapted populations with one-way ANOVA, followed by Tukey’s multiple comparison (42). We compared larval to pupal developmental time, sex ratio, wing size, weight and blood intake using the t-student test (42). We tested differences in longevity using the Log-rank (Mantel Cox) analysis (42). We compared oviposition patterns through a cubic spline analysis (42).

### Crosses between laboratory-adapted populations

We performed reciprocal crosses between mosquitoes of the Tap and Fo populations by exposing 20 newly emerged females of one population to 7 newly emerged males of the other population for seven days. Then, we offered a blood meal and provided a damp filter paper for egg deposition. F_1_ eggs were hatched, and the progeny of each cross was let interbreed. We determined wing length, fecundity and fertility of F_1_ mosquitoes as described above. We compared wing length, fecundity and fertility of the progeny of each cross, also with respect to those of the parental Tap and Fo populations, using a Mann-Whitney test in Prism 9.1 because data were not parametric (GraphPad).

### Energy reserve

We sampled a total of 36, 3-5 days old females before blood meal, 24 and 96 hPBM. We dissected the fat body of each individual in 75% ethanol and weighted it using a microbalance (Mettler AC100). We homogenized samples individually in 180 μL of lysis buffer (100 mM KH_2_PO_4_ (Sigma-Aldrich), 1 mM DTT, 1 mM EDTA (Thermo Fisher) pH 7.4) with pestles. We stored samples at −20°C until processing for quantification of protein, carbohydrate, glycogen and lipid content using a colorimetric assay as previously described (24). Following the same procedure outlined above, we further determined the protein content of the fat body and ovaries of 216 females which were sampled 12, 24, 48, 60 and 72 h pBM. We compared energy reserves and the protein time course with 2-way ANOVA tests followed by Sidak’s multiple comparison tests using Prism 9.1 (GraphPad).

### Midgut trypsin-like activity

We offered a blood meal to 3-5 days old females and sampled fully engorged females after 6, 24, 36, 48 and 60 hours. We also sampled 144 3-5 days old females before BM. We dissected midguts from each mosquito on ice and tested trypsin-like activity as described by Gulia-Nuss et al. (43), with slight modifications. Briefly, we homogenized each sample in 100 μL of an extraction buffer (20 mM Tris/ 20 mM CaCl_2_ [Sigma-Aldrich], pH 8) using a pestle, followed by centrifugation for two minutes at 14000*g*, 4°C. We collected supernatants and stored it at −80°C until measurements. For SF samples and samples collected 6 h pBM we used 10 μL aliquots; for samples of all other time points we used 5 μL aliquots and added them to 100 μL of 4 mM Nα Benzoyl-L-Arginine-p-Nitroanilide (BApNA; Sigma-Aldrich) in a 96-well plate. We incubated plates at 37°C for 10 min, followed by absorbance reading at 405 nm in a CLARIOstar plate reader (BMG Labtech). We quantified trypsin-like activity using a standard absorbance curve generated using 20 μg of Trypsin from Bovine Serum (Sigma-Aldrich) with BApNA standards (8.96, 4.48, 2.24, 1.12, 0.56, 0.28 and 0.14 mM).

### Microscopy analysis of mosquito ovaries

We offered a blood meal to 3-5 days old females and sampled fully engorged females, in groups of 5, after 24, 30, 36, 42, 48, 72 and 96 hours. We dissected ovaries from each female in 75% ethanol and fixed tissues in Canois solution (60% ethanol (Sigma-Aldrich), 30% chloroform (Sigma-Aldrich) and 10% acetic acid (Sigma-Aldrich)) for up to a week. Afterwards, we rinsed samples three times with phosphate buffered saline (PBS) 1x and permeabilized with permeabilization solution (0.2% Triton (Sigma-Aldrich), PBS 1x and 1% BSA (Promega)) for one hour. Finally, we stained the ovaries with 4’, 6-diamidino-2-phenylindole (DAPI) (Thermo Scientific, diluted 1:50 in PBS 1x) for five minutes before mounting samples on the slide using polyvinyl alcohol medium (Sigma-Aldrich). We took images with a TCS Sp8 confocal microscope (Leica) at Centro Grandi Strumenti at University of Pavia (https://cgs.unipv.it/?page_id=84).

### Proteomic analyses of ovaries

We offered a blood meal to 3-5 days old females and, either one or four days later, we sampled 100 fully engorged females to dissect ovaries in 75% ethanol under a dissection stereoscope Leica ZOOM 2000. We pooled ovaries from females of the same population and collected at the same time point pBM, for protein extraction following Geiser et al. (44), with some modifications. Briefly, we homogenized tissues in 500 μL of lysis buffer (7M urea, 2M thiourea, 50 mM HEPES pH 8, 75 mM NaCl, 1 mM EDTA and 1x Halt proteases inhibitors cocktail) with a pestle, then we incubated for 30 min in ice, sonicated by probe in ice and centrifuged samples at 12000 rpm for 10 min at 4°C. We collected supernatant and quantified protein content using the Bradford assay (45). Samples were then run on an SDS-PAGE followed by an in-gel digestion. Briefly, 30μg of proteins of each sample were loaded onto an SDS-PAGE gel (stacking 5%, resolving 12%) and proteins were separated in an electrophoretic run at 200V for 40 minutes. The resulting profile was divided into seven fractions and the gel was cut accordingly. Each gel piece was de-staining by covering it with enough volume of a solution of 100mM ammonium bicarbonate and 50% acetonitrile for fifteen minutes. This washing step was repeated until the gel pieces were completely de-stained. Gel pieces were then dehydrated adding pure acetonitrile and incubating at room temperature. Acetonitrile was later dried at 60°C for five minutes. Then we performed reduction with freshly prepared 10 mM dithiothreitol (DTT) solution and incubation at 37°C for 30 min, followed by alkylation with freshly prepared 55 mM iodoacetamide (Sigma-Aldrich) and incubation for 45 min at 60°C. These samples were washed twice with 200 μL of 100 mM ammonium bicarbonate before a second dehydration with 200 μL of pure acetonitrile. We then performed protein digestion by incubating samples on ice for 30 minutes with 50 μL of 20 ng/μL trypsin in 100 mM ammonium bicarbonate, followed by overnight at 37°C with enough 100 mM ammonium bicarbonate to avoid gel drying. The following day, we treated each gel piece with 100 μL of a solution of 50% acetonitrile/5% formic acid, for fifteen minutes at 37°C to extract proteins. Proteins of fragments coming from the same gel fragment were pooled and dried in a Speed Vac® without exceeding 30°C for 4 hours. Peptides were resuspended in 20 μL of distilled water and 1 μL of formic acid and stored at −20°C until mass spectrometry analysis.

We obtained the proteomic profiles of each sample via liquid chromatography-mass spectrometry (LC/MS) at the Centro Grandi Strumenti of University of Pavia (https://cgs.unipv.it/?page_id=96) using the instrument LC/MS-MS LCQ Fleet (Thermo Scientific). We generated the mass spectra using the SCIEX software (Sciex) and identified proteins using a threshold of at least 1 distinct peptide per protein with a 95% confidence and a personalized database, which comprises all proteins of *Ae. albopictus* (AalbF2 assembly) and the *Ae. aegypti* (AaegL5.3 assembly) as obtained from VectorBase (https://vectorbase.org/vectorbase/app).

### Functional gene annotation and enrichment

Gene Ontology (GO) functional assignment and gene enrichment of protein-coding genes followed the strategy outlined by Lozada-Chavez et al., (46). This process allowed annotation of 69% of the *Ae. albopictus* proteome (https://github.com/naborlozada/Khorramnejad_et_al_2024). Our custom annotation database was used for GO enrichment analysis with clusterProfiler v4.2.2 (47) to identify functional groups that were enriched in proteins identified through LC/MS analyses from ovaries. P-values (p≤0.05) obtained with clusterProfiler were corrected for multiple tests with the Benjamini-Hochberg procedure, and redundant enriched GO terms in each major GO classification were removed with clusterProfiler (‘simplify’ function).

### Genomic analyses

We used whole-genome sequencing (WGS) data of pools of 40 mosquitoes/population. Raw sequencing reads are accessible under the NIH BioProject number PRJNA1080321. We assessed the quality of raw sequencing reads with FastQC (v. 0.12.1) (48), and trimmed adapters using fastp (v. 0.23.2) (49). We aligned the resulting reads to the *Ae. albopictus* genome (AalbF2 assembly, VectorBase rel. 55) (50) using the bwa-mem2 aligner (v. 2.2.1) (51). We calculated genome-wide pairwise fixation index (Fst) and Tajima’s D genetic diversity (Table S3) using grenedalf (v. 0.3.0) (52) with the PoPoolation methods (53). Both metrics were computed by excluding nucleotide positions with base counts below 2 (--filter-sample-min-count 2), coverage lower than 5 (--filter-sample-min-coverage 5), minimum mapping quality lower than 20 (-- sam-min-map-qual 20), minimum base quality score lower than 20 (--sam-min-base-qual 20).

## Acknowledgments

We are grateful to Dr Anthony A. James from the University of California, Irvine and all members of the Bonizzoni’s lab. for fruitful discussion.

