## Supplementary figures and images for "Reproductive resource allocation correlates with successful global invasion of a mosquito species"

### Fig.S1

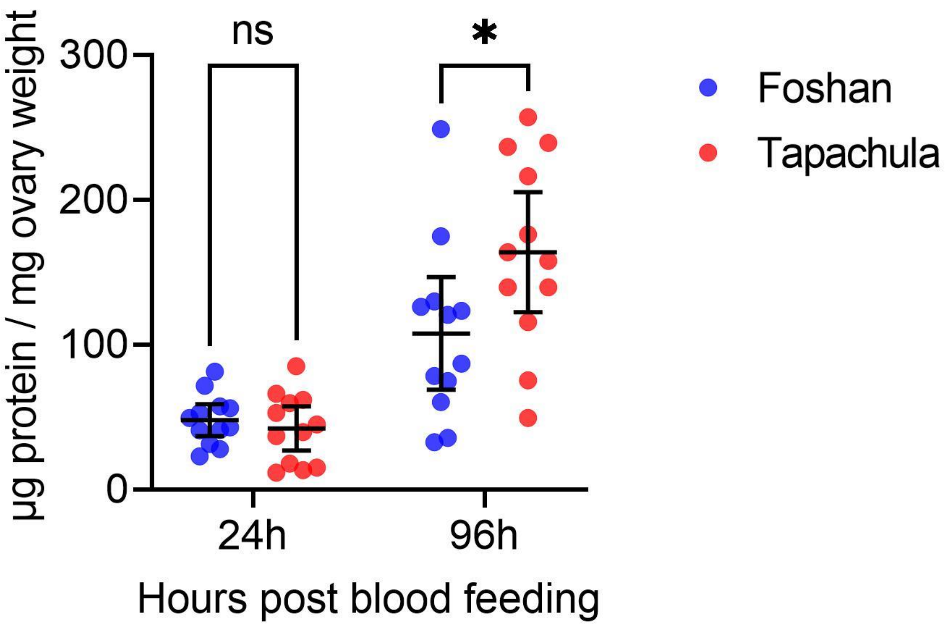

### Fig.S2

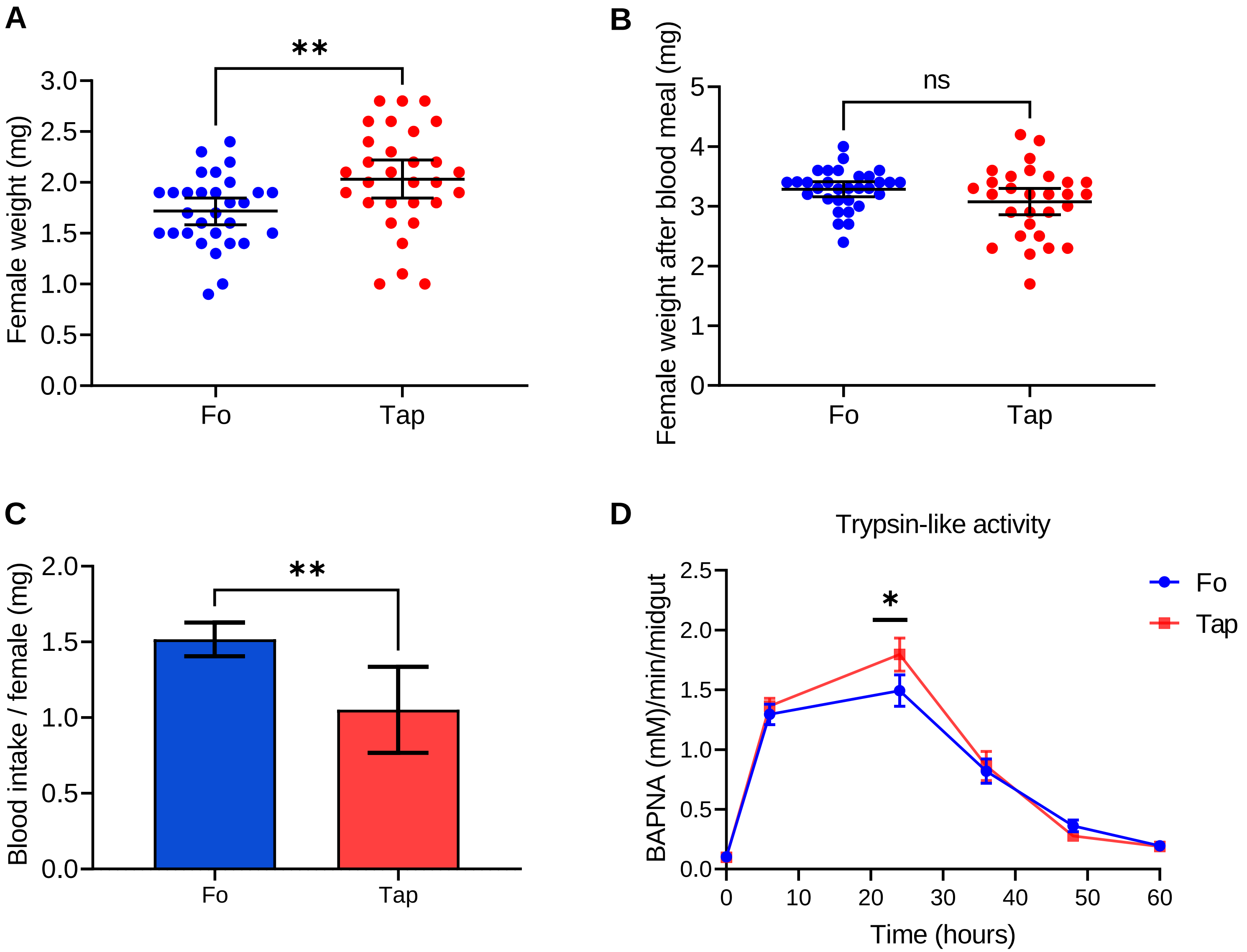
